# A new, Critically Endangered species of *Impatiens* (Balsaminaceae) from the coastal plain of the Republic of Congo

**DOI:** 10.1101/2022.05.05.490806

**Authors:** Martin Cheek

## Abstract

*Impatiens moutsambotei* is described from a herbarium specimen collected at a waterfall in forest on the coastal plain, below the Mayombe Mts of the Republic of Congo. Sharing many distinctive characters with *Impatiens floretii* of the Doudou Mts of Gabon, it is abundantly distinguished inter alia by the leaf-blades which are lanceolate or narrowly elliptic, not ovate, their bases acute, not obtuse or broadly rounded to truncate; the reduced, peduncular bracts bearing long, filiform setae, and not entire; the proximal (upper) of the lateral united petals are entire, not bifid; the spur is curved at the base and overlaps the lower sepal, not curved through its length to describe a semi-circle and held below the lower sepal. Both species are placed in the *Impatiens macroptera* species aggregate. *Impatiens moutsambotei* is known from a single site, where it was collected nearly 30 years ago and not seen since. The new species is assessed as Critically Endangered due to threats of habitat clearance from mining and road widening, and may be already be extinct.

## INTRODUCTION

*Impatiens floretii* Hallé & A.M. Louis of Gabon is one of the most distinct and remarkable African species of the genus (Hallé & Louis 1989). It combines opposite leaves, which are seen in only a minority of the African species, with leaf-teeth extended into setae of more than 10 mm long: longer than in any other African species. It also has unusually large, broad, leafy bracts. The broad lateral sepals extend almost the full width of the pre-anthetic flower bud in profile (in most species they are minute) while the lower sepal-spur has a shovel-like mouth rather than being truncate as in most species. The plant is glabrous apart from the proximal part of the peduncle which is densely patent-puberulent, and the abaxial leaf-blade which is scattered hairy. This species in endemic to the Doudou Mts of southern Gabon where it mainly occurs on granite inselbergs (Sosef et al. 2004).

During January 2020, the author was reviewing unidentified material of *Impatiens* from West-Central Africa in the Kew Herbarium while identifying *Impatiens* specimens for the Cameroon TIPAs programme (Darbyshire et al. 2017), two of which proved to be new species to science (Cheek et al. 2022). Among the unidentified specimens was one, *JJFE de Wilde* 11072 (K) from the Mayombe Mts of the Republic of Congo (Congo-Brazzaville) with all the features described above for *Impatiens floretii*. However, since the specimen was from a location 280 km distant, in a different habitat (a waterfall in lowland forest), indeed in a different country from *I. floretii*, it was studied in detail. Although sharing so many unusual morphological traits with that species, it also showed numerous points of difference (Table 1) below. Therefore, it is here described as a new species, and named as *Impatiens moutsambotei* Cheek.

**Table 1.** Diagnostic character separating *Impatiens floretii* and *Impatiens moutsambotei*. Data for first species from Hallé and Louis (1989), *van Valkenburg* 2718 (K), and *Jongkind* 5782 (K).

|  | *Impatiens floretii* | *Impatiens moutsambotei* |
| --- | --- | --- |
| Height | 0.2-0.6m | to 1.2m |
| Length of longest petioles (on main axis) | 3.8-5cm | 8-10.5cm |
| Leaf-blade shape (length:breadth ratio) | Ovate (<2:1) | Lanceolate or narrowly elliptic (>2:1 to 4.5:1) |
| Leaf base | Obtuse or rounded | Acute |
| Peduncular bracts | Entire, filamentous setae absent | Margins with filamentous setae 4-6.8 mm long |
| Bracts (subtending pedicels) | Flat, transversely elliptic, c. 3 x 4 mm, apex retuse-mucronate. | Sheathing, oblong-elliptic, 7-8 x 5-6 mm, apex acute |
| Flowers per inflorescence | 1 - 2(-3) | 4 - 6 |
| Flower length | 3.5 cm | 1-1.3 cm |
| Lateral sepals | Elliptic | Obovate |
| Spur | Curved throughout length, held below lower sepal | Curved at base, overlapping with lower sepal |
| Proximal petals of lateral united petals | Conspicuously bilobed | Entire |
| Pedicel length | 8-12 mm | 14-15(-17) mm |
| Dorsal sepal | 10-11 x 7 mm with dark transverse bars | c. 8 x 8 mm, transverse bars absent |

*Impatiens* L. (Balsaminaceae) with over 1067 species accepted (Plants of the World Online, continuously updated), is one of the most species-diverse genera of vascular plants, and is nearly cosmopolitan, indigenous species being absent only from S. America and Australia. The current taxonomic framework for continental Africa was established by Grey-Wilson (1980) who recognised 110 species in his monograph. Since then, 24 further species have been published (Abrahamczyk et al. 2016; Bos 1991; Cheek and Csiba 2002; Cheek and Fischer 1999; Cheek et al. 2022; Fischer 1997; Fischer et al. 2003; 2021; Frimodt-Möller and Grey-Wilson 1999; Grey-Wilson 1981; Hallé and Louis 1989; Janssens et al. 2009a; 2010; 2011; 2015; 2018; Pócs 2007). The three broad centres of species diversity in tropical Africa are the western African mountains, primarily the Cameroon Highlands (28 species), the E. Arc mountains of Tanzania with the Kenya Highlands (24 species) and the Albertine Rift (20 species) (Fischer et al. 2021).

Africa was colonised by *Impatiens* from SW China on three occasions (Janssens et al. 2009b). The first colonisation was in the Early Miocene (clade A1, E. and S. Africa). Two further colonisations occurred in the Late Miocene or Early Pliocene, namely clade A2, endemic to W. Africa, and clade A3, which gave rise to the largest diversification in Africa, mostly in E and E-Central Africa, but with some species in both W and C Africa and then generally geographically disjunct between these two locations, e.g, *Impatiens mannii* Hook. f. and *I. burtonii* Hook. f. (Janssens et al. 2009b). Most speciation has occurred in the Pleistocene and has been rapid, e.g, two Gabonese species which are calculated to have diverged only 0.18 million years BP (Janssens et al. 2011). Many of the species are geographically localised, several to individual mountains. In Africa, *Impatiens* characterise humid tropical and subtropical montane forests above 500 – 800 (– 5000) m alt. Most species of *Impatiens* cannot survive drought or extended exposure to direct sunlight. As a result, *Impatiens* species are typically confined to stream margins, waterside boulders, and wet and/or montane forests (Fischer, 2004). The species are pollinated by insects, including bees and butterflies, but with at least ten species which are adapted in multiple ways to being pollinated by sunbirds (Bartoš et al. 2012; Bartoš and Janeček 2014; Hořák and Janeček 2021). Studies of flower structures in relation to insect pollination currently lag behind those for bird pollinated species. Seed dispersal is mediated by explosive fruits (Grey-Wilson 1980).

## MATERIALS AND METHODS

This study is based on herbarium specimens. All specimens cited have been seen unless indicated as “n.v.”. Terms follow Beentje and Cheek (2003) and Grey-Wilson (1980). Herbarium citations follow Index Herbariorum (Thiers et al. continuously updated), nomenclature follows Turland et al. (2018) and binomial authorities follow IPNI (continuously updated). The monograph of African *Impatiens* (Grey-Wilson 1980) was the principal reference work used and the herbarium of RBG, Kew (K) to determine the identification of the specimen of what proved to be a new species. Material of the suspected new species was compared morphologically with protologues, reference herbarium specimens, including type material of W-C African *Impatiens* principally at K, but also using material and online images from BR, MO, P and YA.

Google Earth was used to study the locality remotely with satellite imagery, guided by the map reference on the herbarium specimen. The conservation assessment was made using the categories and criteria of IUCN (2012). Herbarium material was examined with a Leica Wild M8 dissecting binocular microscope fitted with an eyepiece graticule measuring in units of 0.025 mm at maximum magnification. The flowers were rehydrated before being measured and drawn. The drawing was made with the same equipment using a Leica 308700 camera lucida attachment.

## RESULTS AND DISCUSSION

The characters separating *de Wilde et al*. 11072 (K) from *Impatiens floretii* are indicated in Table 1 below. Both species appear to belong to the *Impatiens macroptera* species aggregate of Grey-Wilson (1980), owing to the pedunculate, sub-umbellate racemes, the broadly saccate lower sepal which abruptly constricts into a long, incurved spur, the cucullate dorsal petal, and the large lateral united petals.

***Impatiens moutsambotei*** Cheek **sp. nov**. (Figures 1 and 2)

**Figure 1.**
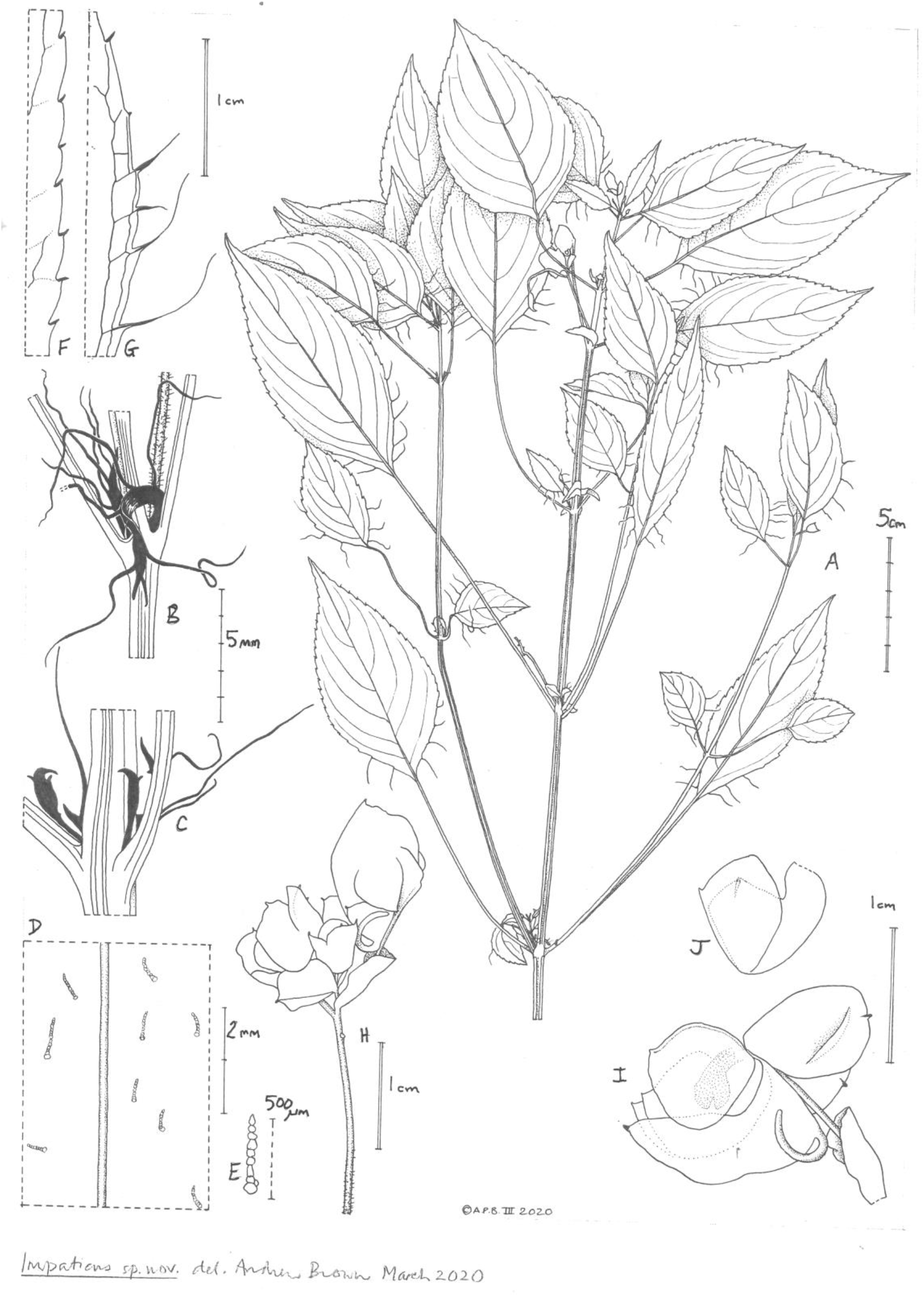
*Impatiens moutsambotei*. A. habit; B. C. nodes with (B) and without (C) hairy peduncle, showing the bract-like axillary leaves (shown in black); D. adaxial surface of leaf-blade near midrib showing sparse hairs; E. detail of moniliform multicellular hair; F. marginal serrations from midlength of leaf-blade; G. marginal serrations with teeth extended into filiform setae from proximal margin of leaf-blade; H. inflorescence with 1 flower fallen; I. flower with lateral sepals raised to show remainder of flower; J. lateral united petals showing the highly overlapping apical and basal petals. Scale bars: A = 5cm; F-J = 1cm; BC = 5 mm; D= 2 mm; E= 500um. All drawn from *de Wilde et al*. 11072 by Andrew Brown.

Type. Congo-Brazzaville, 4km on the road N6 Louvoulou-Kakamoéka. 4°20’S 12°09’E, Alt: 60m. fl. 12 February 1993, *JJFE de Wilde*, with L.J.G. van der Maesen and J.-M. Moutsamboté 11072 (holotype K barcode K000593349; isotypes IEC n.v, WAG n.v.).

### Diagnosis

*Impatiens moutsambotei* differs from *I. floretii* Hallé and J.M. Louis in that the leaf-blades are lanceolate or narrowly elliptic, not ovate, their bases acute, not obtuse or broadly rounded to truncate; the peduncular bracts (at the base of the inflorescences) bear long, filiform setae, and are not entire; the proximal united petals are entire, not bifid; the spur is curved at the base and overlaps the lower sepal not curved through its length to describe a semi-circle and held below the lower sepal; the bract is sheathing, c. 7 × 5 mm the apex acute, not flattened, 3 × 4 mm, the apex retuse-mucronate. Additional diagnostic features appear in Table 1.

### Description

*Terrestrial herb* to 1.2m tall. Leaves and branching opposite, vegetative parts glabrous, upper internodes of principal axis 6-8.5cm long, axillary branches paired, equal at most nodes.

*Leaf-blades* lanceolate or narrowly elliptic, those of the principal axis 8-11 × 2-4.2(−4.5) cm, apex acutely acuminate, base acute, margin serrate, teeth 9-22 on each side (Fig. 1F), proximal 3-5 pair of teeth with filamentous setae, 0.8-1.3cm long (Fig.1 G), lateral nerves 3-6 on each side of the midrib, arising at c. 50° from the midrib, arcing towards the apex, tertiary nerves inconspicuous. Adaxial surface with thinly scattered simple hairs 0.4-0.8 mm long, hairs moniliform, translucent, 9-14 celled (Fig.1DE). *Petioles* 7.5-8.5(−10.5) cm long, lacking glands or appendages.

*Inflorescences* axillary, one per node, often in successive distal nodes, at the base subtended by two unequal, alternate, leaf-like bracts (Fig. 1 BC), leaf-like bracts ligulate, 3.5-7 × 0.6-0.8 mm, apex acute, one of each inflorescence with 3-5 marginal filamentous setae 4-6.8 mm long, the other with teeth 0.1-0.2 mm long. *Peduncle* c. 17 mm long, ascending, the proximal third to half densely patent puberulent with simple moniliform hairs (Fig.1 BH) as in the leaves; rhachis 5-11 mm long, flowers 4-6, alternate, internodes 2-2.5 mm long. Bracts subtending pedicels, completely sheathing and concealing the developing flower until anthesis, oblong-elliptic, 7-8 × 5-6 mm, apex acute, margins entire. Pedicels 14-15(−17) mm long, ascending. *Flower* 1-1.3 cm long, exterior purple-pink, glabrous. *Lateral sepals* 2, before anthesis enveloping most of the flower, obovate, concave, 9 × 6.5 mm, apical mucro thickened, margin entire. *Lower sepal* broadly saccate, scuttle-shaped, 14 mm long, the distal part of the mouth extending half as long again as the proximal part and with an acute thickened apex. (Fig.1I), spur area lacking transverse bars; spur cylindric, curved through c. 180° near attachment, the distal half only slightly curved, 9 mm long, overlapping with lower sepal, apex rounded, pointing towards the androgynoecium. *Dorsal sepal* cucullate, in profile obovate to orbicular, c. 8 × 8 mm, forming a hood over the gynoecium-androecium, apex acutely mucronate, not thickened, crest absent. *Lateral united petals* c. 10 × 10 mm, stipe (claw) c. 3 mm long, proximal petal quadrangular-rounded, 8-9 × 5 mm, entire, deeply separated from and overlapping ± completely with the distal petal, distal petal obovate, 9 × 6 mm, apex mucronate. Androgynoecium curved, c. 9 mm long. Stamens 5, anthers united in a cap over the ovary, anther cap c. 3 × 3 mm. Ovary cylindric-ellipsoid c. 3 × 1 mm. *Mature fruit* not seen.

### Etymology

Named for Professor Jean-Marie Moutsamboté of the Université Marien Ngouabi and of IEC, the Herbier National of the Republic of Congo (IRSEN, ex ORSTOM) for his national leadership of botany, his encouragement of young Congolese botanists, and for his companionship and cheer in the forests of Congo.

### Phenology

Known in flower only in February.

### Distribution and habitat

Republic of Congo, Kouilou Province, known only from the type location. “Humid rocks and banks shaded by trees. Wet place in half-shade. Cascade alongside road. ..60m alt..” from *de Wilde et al*. 11072. In fact, the altitudinal read from Google Earth imagery is in the range 95-130 m rather than 60 m above sea-level.

### Conservation status

The single known site for this species (description and georeference in metadata of the type specimen cited above) is beside the spur road (RN6) that departs from the main national highway (RN1) that connects Brazzaville and Pointe Noire. Starting from Malélé the RN6 serves the major town of Kakamoéka on the Kouilou River. The road is 4-6 km from and parallel with the western N-S boundary of the Biosphere Reserve of Dimonika that extends up into the Mayombe mountain range. The Dimonika Reserve is reported to be under high pressure (http://datazone.birdlife.org/site/factsheet/dimonika-biosphere-reserve-iba-congo downloaded 2 May 2022) and to exist only on paper, with no effective management (Pintea et al. 2009). The georeference for the site for *Impatiens moutsambotei* is presumably taken from a map (when collected in 1993, global positioning systems (GPS) were rarely available to botanists). Probably for this reason, mapped on Google Earth, the point misleadingly lies 1.5 km west of the road on which it is stated to be next to. At many points along and adjacent to the road, areas have been cleared of forest, possibly for extraction of gold which is reported to be a challenge for conservation of habitat within the Dimonika Reserve (Pintea et al. 2009). Artisanal gold extraction often follows natural watercourses. An area c. 500 m × 200 m has been cleared at the point on the road nearest to the georeference of the type specimen, so it is possible that the site, and the species has been lost already.

If the species survives, it is also at major risk from road widening and straightening for two reasons. Firstly, the road is the main access route to the Sounda hydroelectric dam on the Kouilou near Kakamoéka. This installation has been scheduled for upgrading and extension for at least five years for which an improved access road surface will be needed to bring in heavy machinery and secondly, the road has become the main route for Congo’s timber trucks taking Okoumé (*Aucoumea klaineana* Pierre, Burseraceae) logs from the forests of the Chaillu to the port of Pointe Noire.

Using the IUCN (2012) standard the area of occupancy is estimated as 4 km^2^, being the IUCN-preferred cell-size of that dimension. Given the threats of site destruction (above) we here assess the species as Critically Endangered CR B2ab(iii), assuming that the species is not already extinct. It is to be hoped that *Impatiens moutsambotei* will be found at additional sites which will be secure and offer a better opportunity for protection of this remarkable species.

### Discussion notes

Research of refuge *Begonia* species appears to have led to the discovery of *Impatiens moutsambotei*. Jan J.F.E. de Wilde who led the research at Wageningen (WAG) of African *Begonia* that produced over the years in several publications a near comprehensive revision of the *Begonia* taxa of the continent, was the senior collector of the type and only known specimen of *Impatiens moutsambotei*. At the same site and on the same date as he and his team collected this type specimen, they also collected three taxa of refuge *Begonia*, being *Begonia quadrialata* subsp. *quadrialata* var. *quadrialata (J*.*J*.*F*.*E. de Wilde et al*. 11082), *B. quadrialata* subsp. *quadrialata* var. *pilosa* Sosef (J.J.F.E.*de Wilde et al*. 11081, 11092) and *B. potomaphila* Gilg (J.J.F.E. *de Wilde et al*. 11083, 11084). These collections are documented by Jan J.F.E. de Wilde’s former PhD student, Marc Sosef, (Sosef 1994). It is not unusual for species of refuge *Begonia* and of *Impatiens* to grow together at suitable micro-sites in Guinea-Congolian Africa. Both groups thrive in cool, humid, partial-shaded environments that occur at cascades, as at this site. Gesneriaceae and ferns are other groups typically found at cascade sites such as the type location of the new species, together with Podostemaceae (Cheek et al. 2004)

### Biogeographical separation of Mts Doudou of Gabon and the Mayombe Mts

The geographical separation between the sites for *Impatiens floretii* at the Mts Doudou in Gabon, and the site for *Impatiens moutsambotei* on the coastal plain at the foot of the Mayombe Mts of Congo, is about 280km as measured on Google Earth. These two ranges are treated as separate Pleistocene Refuges by Sosef (1994) in his study of *Begonia* sect. Loasibegonia and sect. Scutibegonia (the refuge *Begonia* groups). Although separated by no great geographical distance, Mts Doudou are placed in a separate cluster of refugia from Mayombe due to the disparity between the refuge *Begonia* taxa that they each contain (Sosef 1994). Despite this, there seems little doubt that these two *Impatiens* species are closely related since they share so many unusual morphological characteristics as indicated in the introduction, however convergent evolution cannot be ruled out.

*Impatiens moutsambotei* will be the seventh accepted taxon of *Impatiens*, and the first national endemic of the genus, documented from the Republic of Congo according to Plants of the World online (continuously updated). However, the country is one of the least well-surveyed countries for plant diversity in the tropical African forest zone. In comparison adjacent Gabon has 22 *Impatiens* taxa (Plants of the World Online continuously updated; Sosef et al. 2006) and Cameroon 28 taxa including seven that are confined (endemic) to the Cross-Sanaga Interval (Cheek et al. 2001). The Cross-Sanaga has the highest vascular plant species, and highest generic diversity per degree square in tropical Africa (Barthlott et al. 1996; Dagallier et al. 2020, respectively), including endemic genera such as *Medusandra* Brenan (Peridiscaceae, Breteler et al. 2015; Soltis et al. 2007).

New species to science continue to be published in the Republic of Congo, from small herbs to canopy trees of rainforest. Among the new species published since 2010 are four species of *Aframomum* (Zingiberaceae, Harris and Wortley 2018), *Baphia vili* Cheek (Leguminosae, Cheek et al. 2014), *Campylospermum glaucifolium* Biss. (Ochnaceae, Bissiengou 2013), *Carpolobia gabonica* Breteler (Polygalaceae, Breteler 2010), *Chassalia lutescens* O.Lachenaud and D.J.Harris (Rubiaceae, Lachenaud and Harris 2010), *Craterispermum capitatum* Taedoumg and De Block (Rubiaceae, Taedoumg et al. 2017), two species of *Gilbertiodendron* (Leguminosae, van der Burgt et al. 2015), *Kalaharia schaijesii* Bamps (Labiatae, Bamps 2013), *Morinda mefou* Cheek (Rubiaceae, Cheek et al. 2011), *Octoknema chailluensis* Malécot and Gosline (Olacaceae, Gosline and Malécot 2012), *Salacia arenicola* Gosline (Celastraceae, Gosline et al. 2014), and *Vepris teva* Cheek (Rutaceae, Langat et al. 2021)

## CONCLUSIONS

Many of the new species to science being described today, such as *Impatiens moutsambotei*, are rare and range-restricted, making them likely to be threatened. Other examples of range-restricted species rom those listed above as recently published from the Republic of Congo include *Octoknema chailluensis, Salacia arenicola*, and *Vepris teva* all of which have provisional conservation assessments of threatened. However, there are some exceptions, species which are both new to science, frequent and wide-ranging (e.g. Bamps 2013, Cheek and Etuge 2009; Cheek et al. 2019a). Until species are delimited and known to science, it is much more difficult to assess them for their conservation status and so the possibility of protecting them is reduced (Cheek et al. 2020).

To increase the survival of such newly discovered range-restricted species there is an urgent need to assess the species for their extinction risk, applying the criteria of the IUCN Red List of Threatened Species (Bachman et al. 2019) and publishing them on the official red list iucnredlist.org. The majority of plant species still lack such assessments (Nic Lughadha et al. 2020). Documented extinctions of plant species are increasing (Humphreys et al. 2019) and recent estimates suggest that as many as two fifths of the world’s plant species are now threatened with extinction (Nic Lughadha et al. 2020). Recent examples of species found to be extinct in the Lower Guinea phytogeographical region (W-C Africa) are *Oxygyne triandra* Schltr. and *Afrothismia pachyantha* Schltr. (Cheek and Williams 1999; Cheek et al. 2018a; Cheek et al. 2019b). In some cases, species appear to have become extinct even before they are named for science e.g.in Gabon, the spectacular aroids *Pseudohydrosme bogneri* Cheek and Moxon-Holt (Moxon-Holt and Cheek 2021) and *P. buettneri* Engl. (Cheek et al. 2021) and in Cameroon the cloud forest tree *Vepris bali* Cheek (Cheek et al. 2018b).

It is hoped that discovery, formal publication and future Red Listing of additional threatened endemic species such as *Impatiens moutsambotei* will help support the motivation and case for resources for the protection of the natural areas in which they occur, such as through designation of Tropical Important Plant Areas (TIPAs) (Darbyshire et al. 2017).

## ACKNOWLEDGEMENTS

The discovery of *Impatiens moutsambotei* in the Kew Herbarium was a side product of work on the Cameroon species of *Impatiens* which was supported by funding of the Cameroon TIPAs programme from Players of People’s Postcode Lottery (PPL). Janis Shillito typed the manuscript. Two anonymous reviewers are thanked for constructively reviewing an earlier version of this paper.

The authors declare no conflict of interest.

